# HSP90 buffers newly induced mutations in massively mutated plant lines

**DOI:** 10.1101/355735

**Authors:** G. Alex Mason, Keisha D Carlson, Maximilian O Press, Kerry L Bubb, Christine Queitsch

**Affiliations:** Department of Genome Sciences, University of Washington, Seattle, WA, 98195, USA; Department of Biology, University of Puget Sound, Tacoma, WA, 98416, USA; Phase Genomics Inc., Seattle, WA, 98195

## Abstract

Robustness to both genetic and environmental change is an emergent feature of living systems. Loss of phenotypic robustness can be associated with increased penetrance of genetic variation. In model organisms and in humans, the phenotypic consequences of standing genetic variation can be buffered by the molecular chaperone HSP90. However, it has been argued that HSP90 has the opposite effect on newly introduced genetic variation. To test the buffering effect of HSP90 on new mutations, we introduced vast numbers of mutations into wild-type and HSP90-reduced plants and assessed embryonic lethality and early seedling phenotypes for thousands of offspring. Although the levels of newly introduced mutations were similar in the two backgrounds, the HSP90-reduced plants showed a significantly greater frequency of embryonic lethality and severe phenotypic abnormalities, consistent with higher penetrance and expressivity of newly introduced genetic variation. We further demonstrate that some mutant phenotypes were heritable in an HSP90-dependent manner, and we map candidate HSP90-dependent polymorphisms. Moreover, both sequence and phenotypic analyses of wild-type and HSP90-reduced plants suggest that the HSP90-dependent phenotypes are largely due the newly introduced mutations rather than to an increased mutation rate in HSP90-reduced plants. Taken together, our results support a model in which HSP90 buffers newly introduced mutations, and the phenotypic consequences of such mutations outweigh those of mutations arising *de novo* in response to HSP90 perturbation.

## Introduction

In eukaryotes, the molecular chaperone HSP90 facilitates the folding of diverse protein clients (1). These clients are implicated in nearly all aspects of organismal growth and development, and thus HSP90 is highly connected in genetic networks (1–3). As a heat shock protein and negative regulator of heat shock transcription factors (4), HSP90 is also linked to the external environment and enables environmental signaling through chaperoning receptors and downstream signaling components in many pathways (1, 3, 5, 6).

The interaction of HSP90 with its protein clients affords them additional physicochemical stability, allowing clients to accommodate mutations that may otherwise impact their function (7–11). Indeed, across the mammalian lineage, including humans, genes encoding HSP90 client kinases carry significantly more and more deleterious variation that those encoding non-client kinases (10). The higher evolutionary rate of HSP90 clients mirrors results with other chaperones (8, 9, 12–16), suggesting that protein stabilization generally increases evolutionary rates through relaxed selection on new mutations in clients (7).

Consequently, HSP90 inhibition compromises client protein folding and causes deleterious phenotypic effects, which may be exacerbated by genetic variation residing in the genes encoding these clients. This phenomenon is distinct from HSP90-dependent effects on epigenetic phenomena (17–19) or the frequency of *de novo* mutations (2, 20–25). HSP90 perturbation through mutation, pharmacological inhibition, or increased temperature reveals distinct, strain-specific morphological aberrations and altered quantitative traits in flies, plants, fish, and yeast (26–34). In worms, naturally varying HSP90 levels predict the penetrance of mutations in several signaling proteins (35, 36). In humans, HSP90 enables the function of several mutated oncogenic proteins, resulting in the chaperone serving as a drug target in cancer (6, 37–40).

HSP90 can affect both penetrance (*i.e.* frequency of an aberrant trait associated with a genetic variant) and expressivity (*i.e.* severity of an aberrant trait associated with a genetic variant) (25, 29, 31, 33, 35, 36). The effect of HSP90 on variant penetrance and expressivity is the basis of the capacitor hypothesis which posits that HSP90 allows accumulation of genetic variation, which remains phenotypically silent until released upon stress or targeted HSP90 perturbation. However, two controversies, prompted by recent studies, remain to be resolved, the first concerning the nature of the variants that can be buffered by the chaperone (41), and the second concerning the relative contributions of HSP90-dependent *de novo* variation to phenotype (22, 24).

As to the first, the increased evolutionary rate of HSP90 clients across the mammalian lineage suggests that HSP90-buffered variation tends to be relatively old and remains present in populations in the face of selection (10). It is less clear how HSP90 affects new genetic variants or variant combinations that have not been exposed to selection. To address this question, studies in yeast and plants have studied hundreds of divergent segregant lines generated from crosses of outbred strains; these segregant lines represent new variant combinations that have not been exposed to long-term selection (28, 32, 33). In sum, the results argue that HSP90 buffers such new genetic combinations.

A recent study compared several yeast mutation accumulation lines, each carrying ~4 single nucleotide variants, to yeast strains isolated from natural environments to address whether HSP90 acts similarly on newly arising mutations and older standing variation (41). Focusing on trait variance rather than trait frequencies or trait means, Geiler-Samerotte and colleagues argue that HSP90 fails to confer mutational robustness for alleles that have not experienced selection. In contrast to the earlier studies (28, 32, 33), these authors also report diminished rather than increased phenotypic variance in response to HSP90 inhibition in yeast segregant lines, which they posit casts doubt on current strategies to target HSP90 in cancer therapies.

Thus far, no study has evaluated the effects of HSP90 on very large numbers of newly introduced mutations in an isogenic background, a strategy that should resolve these contradictions. Here, we introduced thousands of random mutations into Col-0 plants, the *Arabidopsis thaliana* reference genotype, and into Col-0 plants with reduced HSP90 levels (42) or treated with an HSP90 inhibitor (29). If HSP90 buffers at least some of the newly introduced mutations, we should, and did, observe a greater frequency and severity of mutant phenotypes in HSP90-reduced plants. The HSP90-responsive phenotypes in mutagenized seedlings commonly resembled those arising in the unmutagenized Col-0 background at much lower frequency (29), consistent with the notion that variants in a limited number of client proteins contribute to HSP90-dependent phenotypes as opposed to invoking complex higher-order epistasis across genetic networks. Broadly, our results support the idea that HSP90 buffers new genetic variation.

Lastly, our study design allowed us to address the controversy whether HSP90-dependent phenotypes are largely a consequence of *de novo* mutations arising in HSP90-reduced lines as recently argued (22). Although HSP90 inhibition increases the frequency of certain mutations (2, 20, 21, 23, 24, 43–48), we find no evidence that the majority of HSP90-dependent phenotypes observed here arises due to HSP90-dependent *de novo* mutations. In sum, our results resolve the two remaining controversies surrounding the HSP90 capacitor hypothesis.

## Results

### HSP90perturbation increases penetrance of embryonic lethal mutations

We used ethyl methanesulfonate (EMS) mutagenesis to generate large numbers of new mutations in the *A. thaliana* reference strain Col-0 and in two Col-0-derived HSP90-RNAi lines, HSP90 RNAi-A1 and HSP90 RNAi-C1 (42) (**Figure 1A**). The construct in the HSP90 RNAi-C1 plants interferes with the function of multiple HSP90 paralogs to produce more severe HSP90-dependent phenotypes than the construct in HSP90 RNAi-A1 plants, which interferes only with the heat-inducible HSP90.1 paralog (42).

**Figure 1:**
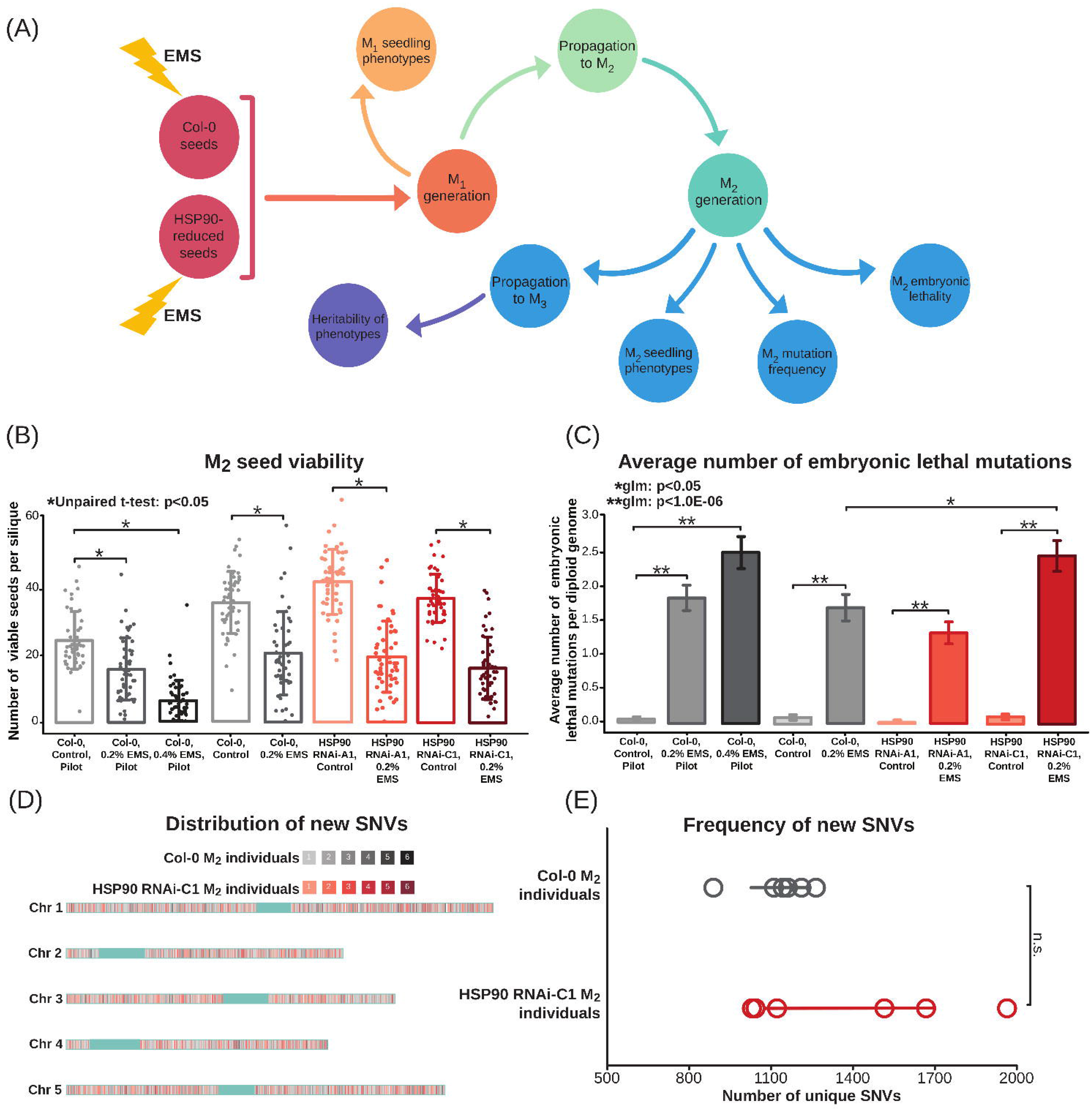
HSP90 perturbation results in greater penetrance of embryonic lethal mutations. (**A**) Experimental workflow across generations. (**B**) Viable M2 seeds per silique after EMS treatment. EMS-treated seeds were grown, yielding M1 plants bearing siliques with M2 seeds. A single silique was collected from 50 individuals for each genotype and viable and dead seeds per silique were counted. *p<0.05, Student’s unpaired t-test. Error bars are standard deviation. (**C**) Estimated number of embryonic lethal mutations per diploid genome (μ) after EMS mutagenesis. μ was calculated as μ=ln(Y)-ln(X), where Y equals the number of siliques collected from M1 plants within a genotype and X equals the number of siliques without dead seeds. *p<0.05, **p<1.0E-06, Poisson regression. Error bars are standard error calculated from a Poisson distribution. (D) Distribution of unique and likely EMS-derived SNVs in six Col-0 M2 and six HSP90 RNAi-C1 M2 individuals. Shades of red denote SNVs detected in HSP90 RNAi-C1 M2 plants, and shades of gray denote SNVs in Col-0 M2 plants. Centromeres were not considered (light blue). (**E**) Frequency of unique and likely EMS-derived SNVs in Col-0 M2 and HSP90-RNAi M2 individuals. Error bars represent 95% confidence intervals. p=0.1460 (not significant), Student’s unpaired t-test.

The EMS-mutagenized seeds (M_1_ generation) were transferred to soil and grown to maturity (**Figure 1A**). To assess the mutagenesis efficiency and the frequency of embryonic lethal mutations, we counted the number of live and dead seeds (M_2_ generation) produced by the M_1_ plants (**Figure 1B, C**) (49). As expected, higher concentrations of EMS led to significantly higher rates of embryonic lethal mutations relative to untreated Col-0 controls (**Figure 1C**). Values for seed viability and embryonic lethal mutations were reproducible across experiments. Untreated plants of the Col-0 and the two HSP90-reduced backgrounds produced similar numbers of live seeds (**Figure 1B**). In contrast, EMS-mutagenized HSP90 RNAi-C1 plants showed significantly more embryonic lethal mutations than EMS-mutagenized Col-0 plants (generalized linear model, family=Poisson: p<1.0E-06) (**Figure 1C**, **S1 Table**). However, EMS-mutagenized HSP90 RNAi-A1 plants did not show more embryonic lethal mutations than EMS-mutagenized Col-0 plants (**Figure 1C**).

A trivial explanation for the greater number of embryonic lethal mutations in the HSP90 RNAi-C1 background is either that these plants were more deeply mutagenized than Col-0 plants or that fewer mutations were repaired. HSP90 chaperones proteins that function in DNA repair (50, 51), and as HSP90 inhibition also increases the rate of certain mutations (2, 20–23, 43–48), even equivalent EMS treatments may produce more mutations in the HSP90 RNAi-C1 background. Alternatively, as predicted under the capacitor hypothesis (25, 31, 52–54), some EMS-generated mutations may acquire greater penetrance when HSP90 is inhibited and therefore result in embryonic lethality.

To distinguish whether increased penetrance or mutation quantity causes the increased embryonic lethality in the HSP90-reduced line, we performed whole-genome sequencing of six randomly selected EMS-mutagenized M_2_ HSP90 RNAi-C1 and six randomly selected M_2_ Col-0 plants. Both backgrounds showed similar genomic distributions and numbers of unique mutations per individual plant (**Figure 1D, E**, **S1 Fig**). Each M_2_ individual showed ~1000 high-confidence, private single nucleotide variants (**Figure 1E**, **S1 Table**, x-coverage >= 5, derived allele coverage >= 2). We conclude that the significantly greater number of embryonic lethals observed in the EMS-mutagenized HSP90-reduced M_2_ progeny are likely due to the increased penetrance of the newly introduced mutations because of reduced HSP90 buffering.

### HSP90 perturbation increases the frequency of severely aberrant phenotypes across generations

Having established that newly introduced mutations are more likely to cause lethality in HSP90-reduced plants than in controls, we examined the effects of HSP90 reduction on the frequency of aberrant morphological phenotypes. As such plant phenotypes are qualitative in nature (29), we used a mutant scoring index to estimate the frequency of 16 complex early seedling traits, categorized by severity (**Figure 2A, B**).

**Figure 2:**
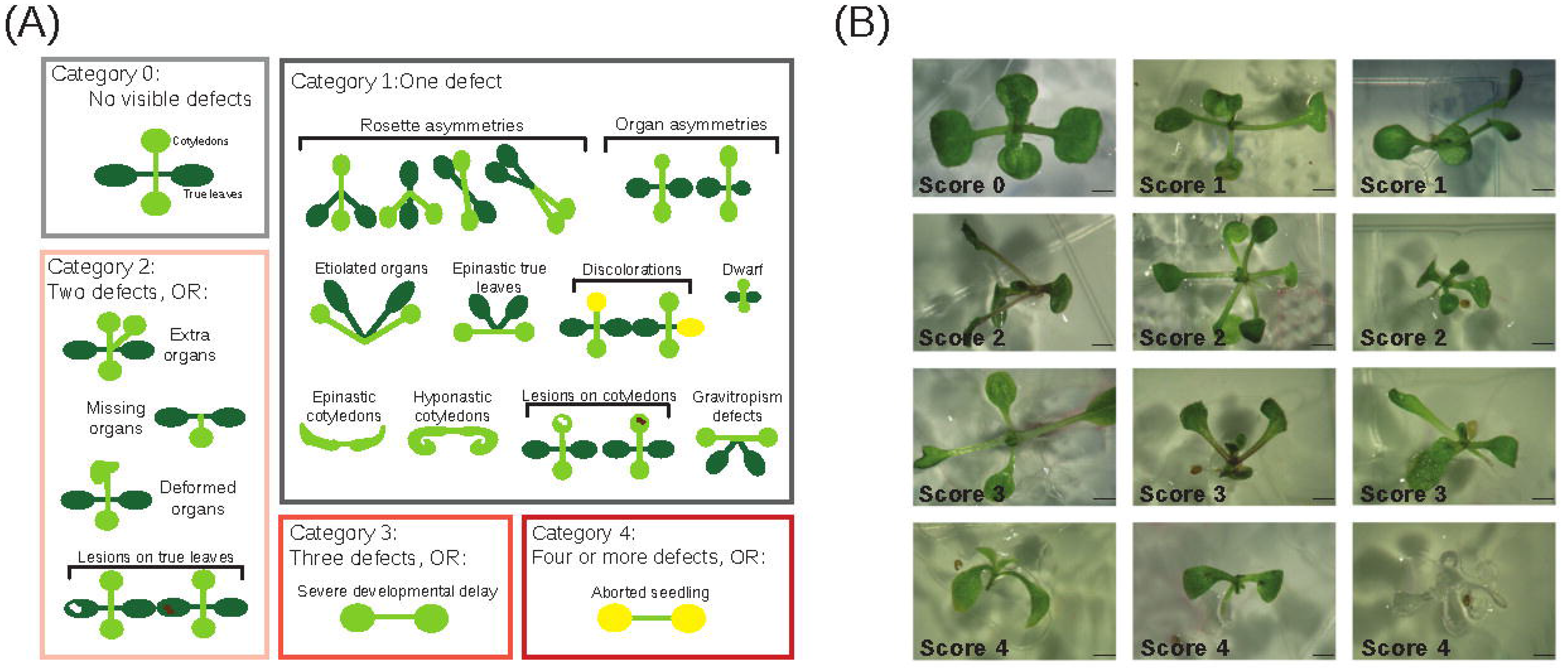
Phenotype scoring index for M1 and M2 seedlings. (**A**) Phenotype categories for 16 aberrant morphological traits in early seedlings, ranging from wild-type-like (category 0) to strongly affected, being either an aborted seedling or showing four or more morphological defects (category 4). (**B**) Example M2 seedlings with common aberrant morphological phenotypes at varying levels of severity. Scores for phenotypic categories are indicated. Scale bars equal 1 mm.

We first examined the frequency of these aberrant phenotypes in the M_1_ plants, in which the EMS-induced mutations are typically heterozygous and sometimes occur only somatically. Heterozygous mutations can show increased penetrance and cause dominant mutant phenotypes when HSP90 is inhibited (55). Indeed, M_1_ HSP90 RNAi-C1 seedlings, but not M_1_ HSP90 RNAi-A1 seedlings, displayed a modest yet significant increase in the frequency of individuals with mutant index scores ≥ 2 (Fisher’s exact test: p-value<0.05, OR=1.5) (**S2 Fig**).

We used the highly specific HSP90 inhibitor geldanamycin (GdA) (Whitesell, 1994) to examine whether a similar trend held for the EMS-mutagenized Col-0 M_1_ plants when HSP90 was inhibited pharmacologically. Indeed, GdA treatment significantly increased phenotype frequency and severity in this population (**S2 Fig**). We conclude that HSP90 perturbation increases the penetrance and expressivity of heterozygous or somatic mutations that affect seedling morphology phenotypes.

Second, we tested the response to HSP90 perturbation, both pharmacologically and genetically, in the M_2_ generation using the same mutant phenotype index (**Figure 3A**). In the M_2_ generation, with the newly introduced mutations segregating, some will be homozygous and others heterozygous. HSP90 perturbation should increase the expressivity and penetrance of these mutations and thereby cause a higher frequency of severely aberrant morphological phenotypes. Consistent with this expectation, we observed a significantly greater frequency of severe phenotypes among the EMS-mutagenized seedlings grown with the HSP90 inhibitor compared to unmutagenized seedlings and mutagenized seedlings grown with the DMSO solvent (**Figure 3B**). This observation manifests statistically as an interaction between GdA treatment and mutagenesis in their effect on phenotype frequency; the strongest such interaction was observed at an intermediate GdA dose (Fisher’s exact test: p-value <1.0E-15, OR=4.6). Genetic HSP90 perturbation yielded similar results: the M_2_ HSP90 RNAi-C1 seedlings produced significantly more aberrant phenotypes than comparable M_2_ Col-0 seedlings (Fisher’s exact test: p-value <1.0E-15, OR=8.8) (**Figure 3C**). We conclude that HSP90 perturbation increases the frequency and severity of aberrant phenotypes after mutagenesis.

**Figure 3:**
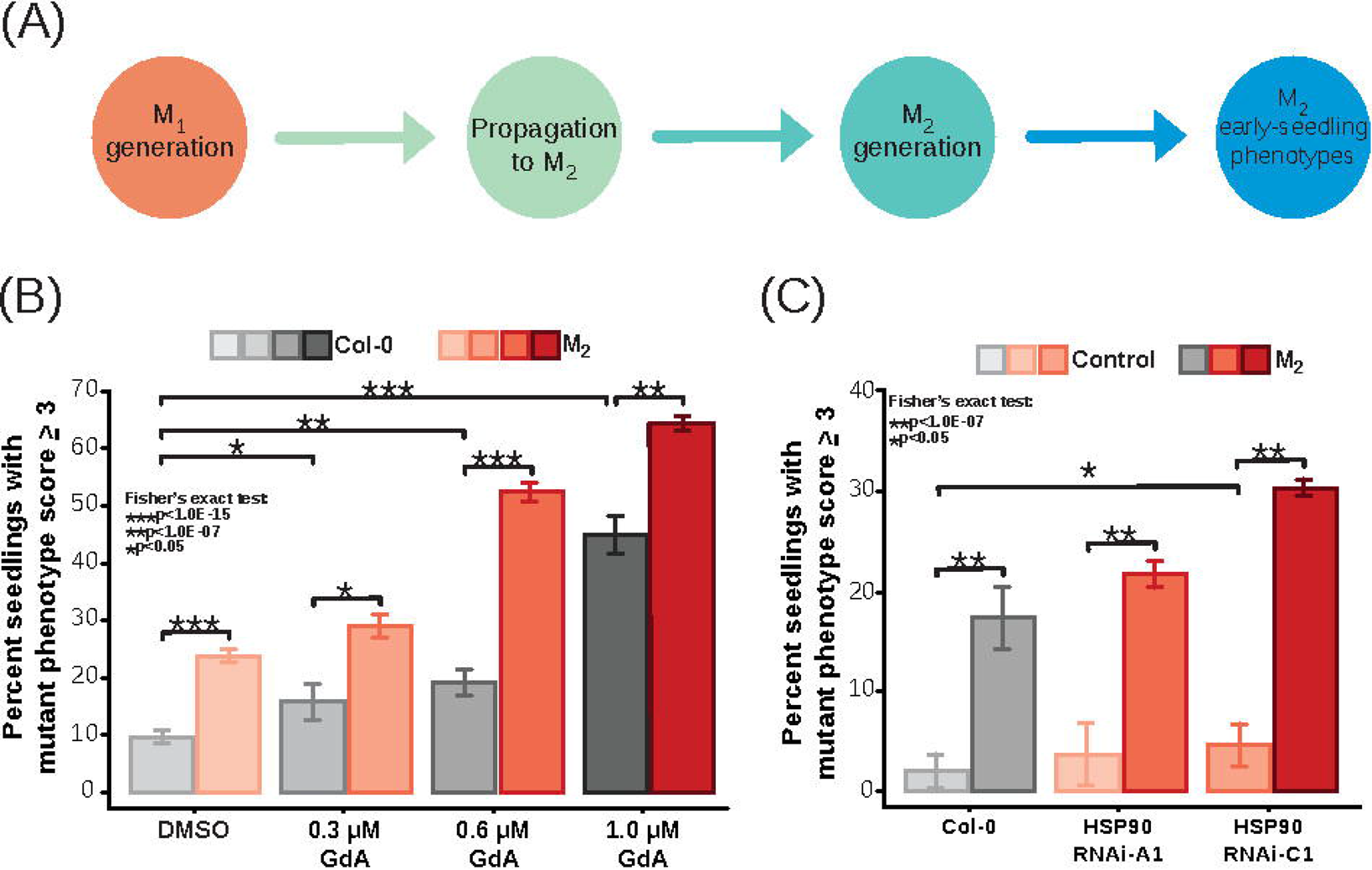
HSP90 perturbation increases the frequency of severely aberrant phenotypes in the M2 generation. (**A**) Experimental workflow with examined seedling populations. (**B**) Phenotype frequencies in response to different concentrations of the HSP90 inhibitor geldanamycin (GdA) in wild-type Col-0 (shades of grey) and mutagenized M2 seedlings (shades of red). Y-axis, percent of seedlings with mutant phenotype scores > 3 (see **Figure 2**). The following numbers of seedlings was examined for each genotype and treatment: wild-type Col-0: DMSO n=731, 0.3 μM GdA n=143, 0.6 μM GdA n=319, 1.0 μM GdA n=243; mutagenized M2: DMSO n=1394, 0.3 μM GdA n=492, 0.6 μM GdA n=933, 1.0 μM GdA n=1488. *p-adj<0.05, **p-adj<1.0E-07, ***p-adj<1.0E-15, Fisher’s exact test. See **S2 Table** for p-values. (**C**) Genetic perturbation of HSP90 reproduces the effects of pharmacological inhibition shown in **B**. Unmutagenized in light shades; mutagenized in darker shades. The following number of seedlings was examined for each genotype and treatment: wild-type Col-0 control, n=347; mutagenized Col-0 M2, n=817. Unmutagenized HSP90 RNAi-A1 control, n=314; mutagenized M2 HSP90 RNAi-A1, n=688. Unmutagenized HSP90 RNAi-C1 control, n=362; mutagenized M2 HSP90 RNAi-C1, n=632. *p-adj<0.05, **p-adj<1.0E-07, Fisher’s exact test. Error bars represent standard error as calculated by: SE = sqrt(p(100-p)/n), where p equals the percentage of seedlings with aberrant phenotypes and n equals the number of observations.

### Tracking the HSP90-dependent nature of aberrant morphological phenotypes across generations

We asked whether the phenotypes from newly introduced mutations are indeed HSP90-dependent, *i.e.* whether or not they would be expressed when HSP90 is fully functional. As it is not possible to examine a single plant both with and without HSP90 perturbation, we generated M_3_ offspring from M_2_ parents that displayed aberrant phenotypes under HSP90-reduced or control conditions (**see Materials and Methods**). Although some of the newly generated EMS mutations will segregate in the M_3_ offspring, this approach generates groups of sibling plants with a similar mutation background that can be examined under either control or HSP90-reduced conditions. Specifically, we selected M_2_ plants with the deformed aerial organ phenotype on DMSO medium (DM plants, Figure 4 A, B) or GdA medium (GM plants). Deformed aerial organs are readily identified and can be a consequence of HSP90 perturbation (29, 42). In addition to the deformed aerial organ phenotype, we also examined the M_3_ progeny of the 22 GM and 10 DM plants for the other 15 phenotypes (**Figure 2A, B**) because more than one phenotype was noted for many parental M_2_ plants (**S3 Table**).

**Figure 4:**
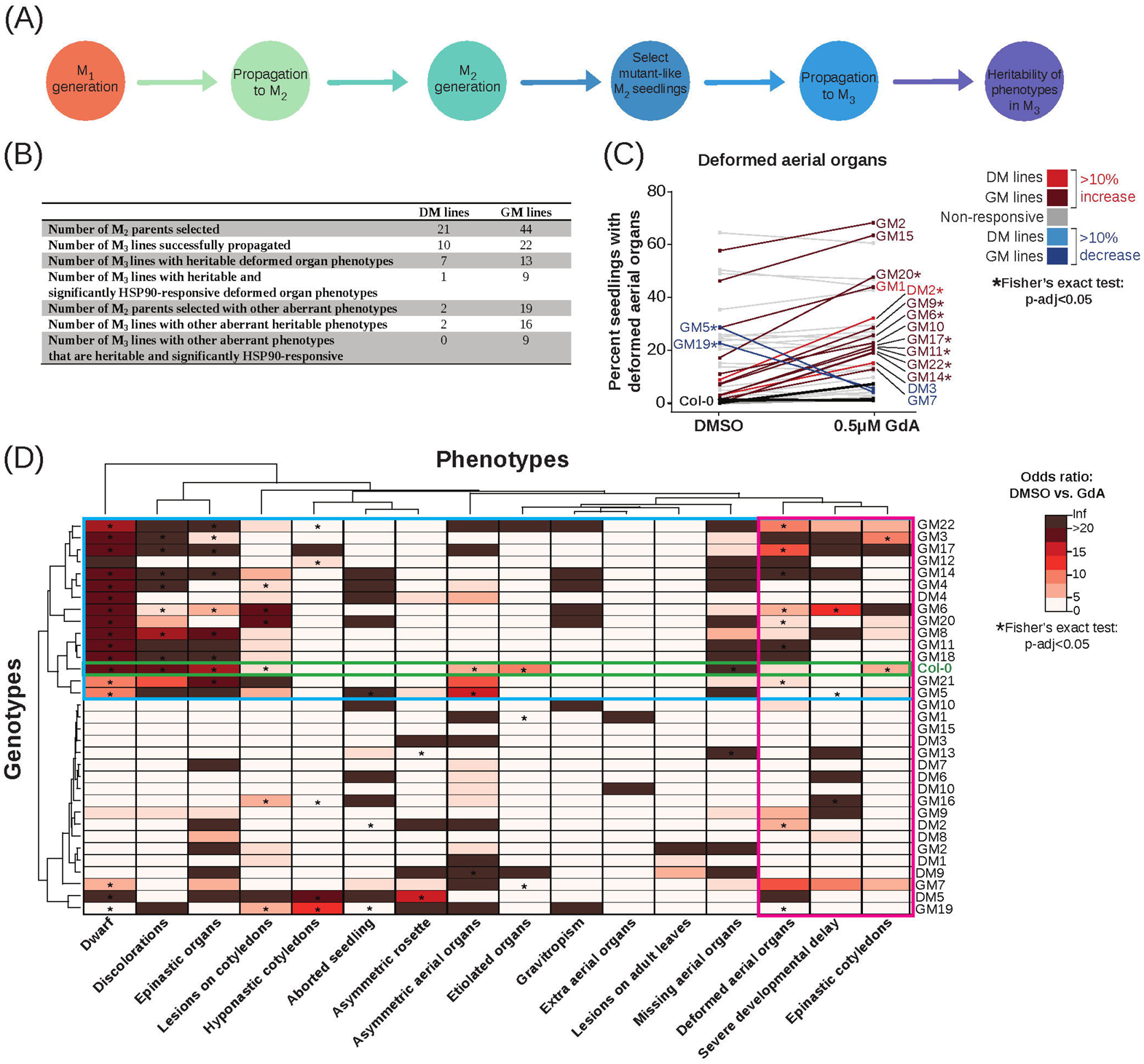
Aberrant morphological phenotypes are heritable and HSP90-responsive across generations in several mutagenized lines. (**A**) Experimental work flow with examined seedling populations. (**B**) Summary table of phenotype heritability in M3 lines. (**C**) Frequencies of the deformed aerial organ phenotype in wild-type Col-0 and mutagenized M3 populations in response to HSP90-reduced (0.5 μM GdA) or mock growth conditions (DMSO). Black lines represent the response of wild-type Col-0 to HSP90 perturbation in three biological replicates. Shades of red denote lines with 10% (5% for frequency of aborted seedlings) increase in phenotype frequency under HSP90-reduced conditions and shades of blue denote lines with 10% decrease in phenotype frequency under HSP90-reduced conditions (5% for frequency of aborted seedlings). Gray line color denotes non-responsive lines. The following number of seedlings was examined: n~144 for wild-type Col-0 and n~72 for each M3 line. *p-adj<0.05, Fisher’s exact test. *p-adj<0.05, Fisher’s exact test. (**D**) Heatmap of clustered odds ratios for GdA vs. DMSO (mock control) comparisons as calculated for wild-type Col-0 (green outline) and the mutagenized M3 lines across 16 early-seedling phenotypes (see **S2 Table** for all p-values and odds ratios (OR), **Figure 2**). Blue outline indicates M3 lines showing HSP90-responsive phenotypes commonly observed in Col-0 wild-type seedlings upon HSP90 perturbation albeit at lesser frequency (smaller OR) and severity. *p-adj<0.05, Fisher’s exact test.

Under standard growth conditions, 20 M_3_ lines showed significantly increased frequencies of deformed aerial organs, with more DM lines (7 out of 10) than GM lines (13 out of 22) producing this phenotype (**Figure 4A, B**). The lesser heritability of this trait in the GM lines is consistent with HSP90 perturbation increasing penetrance and expressivity of newly introduced variation in the parental lines.

Next, we asked how many of these lines showed an HSP90-dependent response in the frequency of this phenotype. Nine of 13 GM lines showed significant HSP90-dependence in the expression of the aberrant phenotype, whereas only one out of seven DM lines did. This DM line and seven out of the nine significantly HSP90-responsive GM lines showed low frequencies of deformed aerial organs in control conditions (*i.e.* DMSO) and significantly increased phenotype frequencies when HSP90 was perturbed (**Figure 4C**, GM, dark red lines, DM, bright red line). This HSP90-dependent increase in phenotype frequencies agrees with several previous studies (29, 30, 34, 36, 56). We also found two GM lines with a significantly decreased frequency of the aberrant phenotype (**Figure 4C**, GM dark blue lines, Fisher’s exact test: p-adj<0.05, OR<0.03), reminiscent of previous findings that HSP90 perturbation can conceal the phenotypic consequences of underlying variation (26, 28, 32, 41).

As some parental M2 plants showed other aberrant phenotypes in addition to the selected phenotype, it was not surprising to find these phenotypes also in their offspring (**Figure 4B**). Only two DM M_2_ plants with deformed aerial organs showed additional defects; both produced M_3_ offspring with these defects; none were HSP90-responsive. In stark contrast, 19 GM M_2_ showed additional aberrant phenotypes; 16 of these produced M3 offspring with these defects; significant response to HSP90 perturbation was observed in nine M_3_ lines (**S3 Fig**).

Assessing all aberrant phenotypes together in the context of genetic background and response to HSP90 perturbation (**Figure 4D**; **S3 Fig; S4 Fig**) revealed important trends. First, consistent with previous data (29, 56), upon HSP90-perturbation, unmutagenized Col-0 wild-type plants tended to show dwarf phenotypes, lesions on cotyledons, discoloration, and epinastic organs among others. Second, about half of the mutagenized M_3_ also showed these phenotypes in response to HSP90 perturbation, albeit often at far higher frequency. Assuming that the HSP90 dependent phenotypes observed in wild-type Col-0 plants reflect dysfunctional client proteins (25, 57), this observation suggests that some of the newly introduced HSP90-dependent variants reside in genes encoding these clients or network nodes closely associated with them. Third, selection for the deformed aerial organ trait in the mutagenized lines increased the phenotype frequencies dramatically in some lines, consistent with a genetic, possibly multigenic basis. Lastly, certain traits, such as aborted seedlings and severe developmental delay that are observed only infrequently in unmutagenized Col-0, were strongly responsive to HSP90 perturbation in the mutagenized M_3_ lines, indicative of problems in early plant development and consistent with the increased embryonic lethality observed in the M_1_ generation. Taken together, our results indicate that at least some newly introduced mutations readily lead to heritable HSP90-responsive phenotypes.

### Mapping HSP90-responsive loci containing newly introduced mutations

To further explore the genetic basis of the HSP90-responsive deformed aerial organ trait, we applied a bulk segregant approach, notwithstanding the significant challenges due to the large number of introduced EMS mutations and the partially penetrant, HSP90-responsive nature of the trait. As the aberrant phenotype was both partially penetrant in the mutagenized lines and present at low levels in Col-0 wild-type seedlings, we would not expect a derived allele frequency of 100% indicating the approximate location of the causal mutation, as expected in a traditional bulk segregant analysis (58). Rather, we expected a contiguous increase in derived allele frequency (DAF) above background levels expected in an F_2_ population in Hardy-Weinberg equilibrium (DAF=0.5).

We selected two M_3_ lines, GM6 and GM9, with significantly increased penetrance of this phenotype under HSP90-reduced conditions (OR>4.5). Single GM6 and GM9 M_3_ individuals with deformed aerial organs were backcrossed to Col-0 wild-type plants to generate heterozygous F_1_ populations. A randomly selected individual from each cross was selfed to generate segregating F_2_ populations for both backgrounds (**Figure 5A, B**; **S5 Fig**). As this backcross will dilute phenotype-associated variation, fewer seedling with deformed aerial organs were observed in standard growth conditions while phenotype frequencies in HSP90-reduced conditions stayed high (**Figure 5C**; **S5 Fig**). To map the HSP90-responsive EMS mutations in the GM6 and GM9 backgrounds, we sequenced pools of approximately 120 F_2_ individuals displaying deformed aerial organs upon HSP90 perturbation. For both backgrounds, we found regions with elevated derived allele frequency, consistent with the presence of newly introduced HSP90-responsive mutations (**Figure 5D**; **S5 Fig; S6 Fig**).

**Figure 5:**
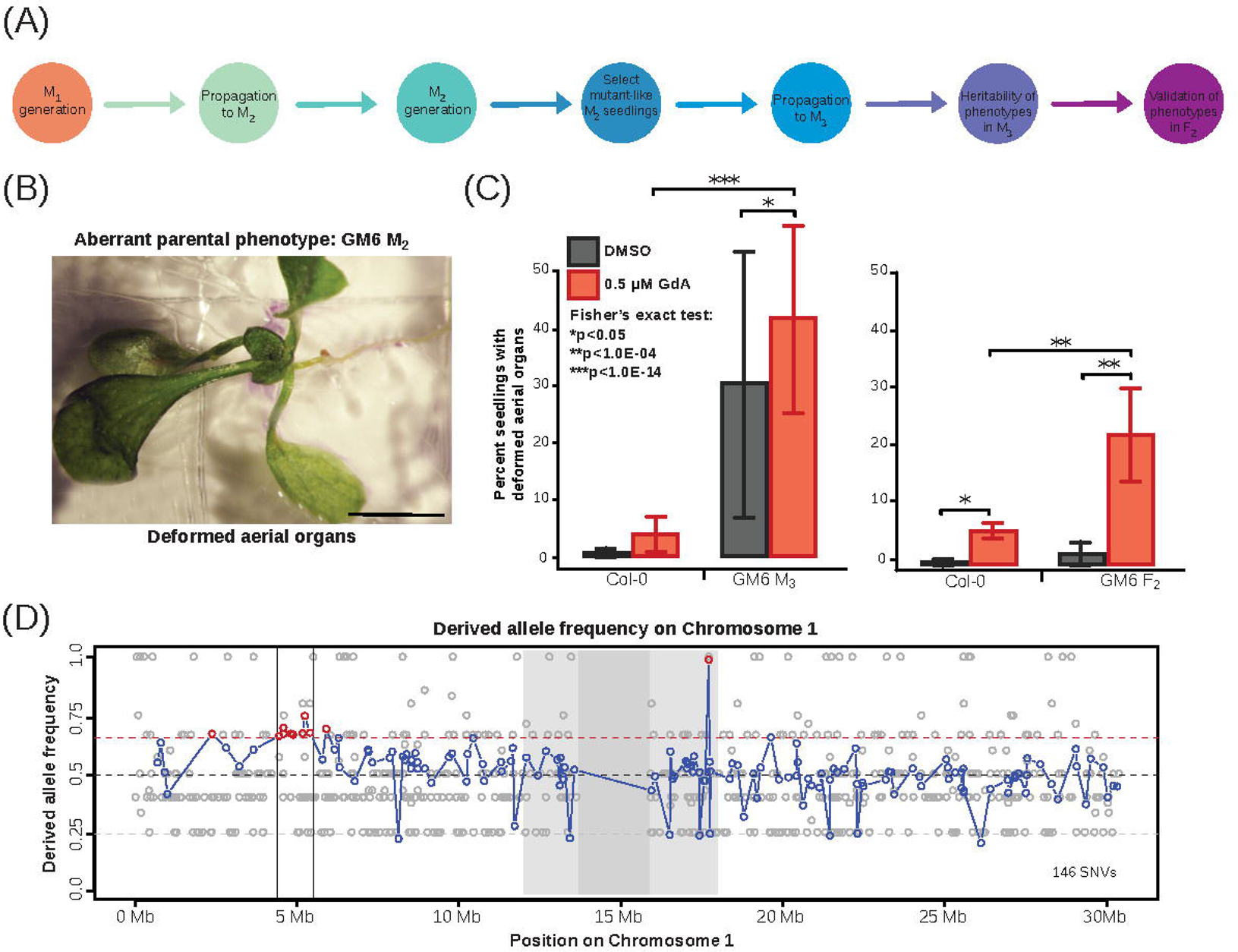
Bulk segregant analysis identifies candidate locus associated with HSP90-responsive trait. (**A**) Schematic of experimental work flow indicating provenance of examined seedling populations. (**B**) Image of the selected M2 parent GM6, showing the deformed aerial organ phenotype that was successfully propagated to the M3 generation. Scale bar equals 5mm. (**C**) The GM6 M3 line showed increased frequency of seedlings with deformed aerial organs compared to wild-type Col-0. A GM6 M3 individual with deformed aerial organs was then backcrossed to wild-type Col-0. The resulting F2 population showed significantly increased frequency of the deformed organs trait when HSP90 was inhibited with GdA. Errors bars represent standard error of the mean across two biological replicates. *p-adj<0.05, **p-adj<1.0E-4, ***p-adj<1.0E-14, Fisher’s exact test. (**D**) Bulk segregant analysis of GM6 F2 populations identifies a candidate locus associated with the HSP90-responsive trait deformed aerial organs. F2 individuals displaying this trait on HSP90-reduced conditions (n=94) were sequenced. Derived allele frequencies (y-axis) were estimated as the fraction of reads supporting the derived allele divided by the number of reads mapping to a given locus (see Materials and Methods). Dark gray box indicates the centromere and light gray box represents pericentromeric regions (chr1: 12,000,000-18,000,000; chr2:1,000,000-7,500,000; chr3:11,000,00-17,000,000; chr4:1,800,000-7,500,000; chr5:10,000,000-17,000,000). Dashed red line represents derived allele frequency of 0.675. Dashed black line represents expected derived allele frequency of 0.5. The candidate locus contains one SNV (chr1:5,407,084 C=>T) encoding a missense mutation (proline to leucine) in the MYB family transcription factor AT1G15720.

For the GM6 F_2_ population, one notable region contained eight consecutive new mutations with elevated derived allele frequency. Six of these did not reside in coding regions, but one (Chr1:5,407,084 C=>T) was a missense mutation (proline to leucine) in the MYB family transcription factor AT1G15720 (**Figure 5D**, **S4 Table**). For the GM9 F_2_ population, one region contained six consecutive new mutations with elevated derived allele frequency. Five of these resided in coding regions, each in different genes, and each was a missense mutation (**S5 Fig**), two of which had negative blosum62 scores, suggesting deleterious effects (**S4 Table**).

To determine how likely these observations were to occur by chance (*i.e*. no new mutation conferred increased phenotypic penetrance upon HSP90-reduction), we performed simulations in which 240 gametes (representing 120 random F_2_ individuals, carrying two haploid pseudochromosomes each) were generated from a simulated F1 individual, heterozygous for *n* mutations, assuming one recombination event per chromosome. Of these 240 pseudochromosomes, we sampled x, reflecting actual experimental x-coverage (read depth, in GM6 mean x-coverage =77; in GM9 mean x-coverage =54). Out of 100 such simulations, none resulted in 5 or more consecutive SNVs with a derived allele frequency greater than 0.65. (**S7 Fig**). Thus, despite the challenges of mapping the genetic determinants of a likely multigenic, partially penetrant trait, our results provide additional evidence for the existence of newly introduced HSP90-buffered variants and support for candidate loci containing them.

### Comparison of the derived allele frequencies in unmutagenized Col-0 wild-type and HSP90-reduced lines

Our study design and sequencing analyses lend themselves to addressing another controversy surrounding the capacitor hypothesis (22): are HSP90-dependent phenotypes largely a consequence of increased frequency of *de novo* germline mutations in response to HSP90 perturbation?

Our Col-0 wild-type (provided in 1994 by Brian Keith) and the HSP90 RNAi lines derived from it have been bulk-propagated in the laboratory for two decades. Therefore, these backgrounds are expected to differ in genome sequence from the *A. thaliana* reference genome. A mutation associated with altered response to HSP90 perturbation would result in an increase in the derived allele frequency for an entire region (haplotype block) containing multiple SNVs, as we describe above for GM6 and GM9. Therefore, we were conservative in annotating SNVs as likely EMS-derived and removed any SNVs that were present pre-mutagenesis in our Col-0 and HSP90-reduced lines compared to the reference genome. To do so, we generated sequence data for Col-0 and the HSP90-RNAi C1 line, examining the number of derived alleles (compared to the *A. thaliana* reference genome) in single plants of both backgrounds (two Col-0 plants, two HSP90 RNAi-C1 plants). As single individuals were sequenced to ~10x coverage, we counted bases only with derived allele frequency of 1 (homozygous for derived allele). We found similar numbers of derived alleles in all four plants (**S1 Table**). There was no evidence supporting higher numbers of single nucleotide mutations in HSP90 RNAi-C1 compared to wild-type Col-0.

Although HSP90 perturbation can lead to *de novo* germline (and likely somatic) mutations, it appears unlikely that these mutations contributed significantly to the greater penetrance and expressivity of aberrant phenotypes in the HSP90-reduced seedlings. This assertion is further supported by the fact that the unmutagenized HSP90 RNAi lines showed similar levels of embryonic lethality as Col-0 wild-type (**Figure 1 B, C**). Moreover, we reproduced the results generated with the HSP90 RNAi-C1 line with pharmacological inhibition of HSP90 in mutagenized Col-0 seedlings. The HSP90 inhibitor is supplied to the growth medium and taken up during seed germination and growth; the seeds contain fully formed embryos. An HSP90-dependent *de novo* mutation affecting these embryos would be somatic and hence unlikely to be heritable. Taken together, our results support the argument that the increased frequency and severity of aberrant phenotypes in response to HSP90 perturbation is due to increased penetrance and expressivity of newly introduced EMS mutations rather than germline *de novo* mutations arising in the HSP90 RNAi-C1 line.

## Discussion

Our data argue that HSP90 buffers the penetrance and expressivity of newly introduced genetic variation in *A. thaliana*. Penetrance effects included an increased frequency of embryonic lethals, and expressivity effects included more severe aberrant morphological phenotypes. Tracking phenotypes across generations, we found several lines that showed heritable aberrant morphological phenotypes in an HSP90-dependent manner. As it does for standing genetic variation (25, 28–31, 34, 59–61), HSP90 plays a role in buffering phenotype against newly introduced genetic variation, which has not been exposed to selection.

These observations are broadly consistent with past studies on new genetic combinations in yeast and plants (28, 32, 33). However, one recent yeast study (41) concludes that phenotypic variance associated with newly introduced mutations typically decreases under HSP90 inhibition (*i.e.* “potentiation”, 25, 26, 62). Here, we argue the opposite: the more parsimonious interpretation of the available data is that HSP90 commonly buffers new mutations.

Our argument relies on multiple lines of evidence. First, we screened a large number of mutations. The mutagenized and sequenced M_2_ individuals carried between 800 to 2,000 new mutations, such that we screened on the order of tens of millions of new mutations, many of them through multiple generations. In contrast, the 94 mutation accumulation lines tested in the recent yeast study carry on average only four single nucleotide mutations per haploid genome (41). Second, we demonstrated the substantial heritability of HSP90-responsive traits, suggesting that these phenotypes are likely to be the effect of genetic mutations rather than stochastic effects. Third, we identified candidate loci associated with an HSP90-responsive phenotype in two independent lines. Fourth, we used two methods (genetic and pharmacological) to perturb HSP90, whose results corroborated each other, demonstrating that the findings are not a consequence of either geldanamycin’s off-target effects or a peculiarity of the HSP90 RNAi lines. Fifth, all mutations were assayed immediately after introduction into the *A. thaliana* genome, such that our analysis should represent the full spectrum of phenotypic effects of new mutations, including those that would be quickly purged by selection. Last, in prior studies, we demonstrate that the effects of revealed HSP90-responsive variation on phenotype means outweigh its effects on phenotype variance by as much as a magnitude, consistent with the capacitor hypothesis’ premise that HSP90-dependent phenotypes can be acted upon by natural selection (33).

However, the present study has some limitations. First, the semi-quantitative phenotype categories make estimates of heritability subject to threshold effects, likely decreasing heritability estimates. Second, aside from embryo lethality, we assayed highly complex developmental phenotypes, and the effects of HSP90 on these phenotypes may be less relevant in the unicellular yeast, which does not experience a prolonged ontogeny. Third, EMS mutagenesis may introduce a different spectrum of mutations than those that arise spontaneously in natural populations.

That HSP90 can buffer new mutations is also supported by newly emerging data. For example, saturation mutagenesis of the non-client yeast transcription factor Ste12 revealed HSP90-responsive Ste12 variants; these variants were rare and position-dependent; moreover, they were implicated in the environmentally-sensitive balance between the mutually exclusive traits of yeast mating and invasion that are governed by Ste12 (63). Similarly, recurrent HSP90-dependent variants in oncogenes are highly sequence-specific and rare (60, 64–66). Notwithstanding these results on the buffering of new mutations, standing variation is likely a more significant actor in most HSP90-dependent phenotypes (11, 25–34, 56, 59).

## Materials and Methods

### Plant Growth Conditions

Seedlings of the Columbia-0 (Col-0) ecotype were used as a wild-type reference; this stock has been propagated in the Lindquist and Queitsch labs since 1994 and was originally provided by Brian Keith. The HSP90 RNAi-A1 and HSP90 RNAi-C1 were generated in this background (42). For experiments, seeds were sterilized with ethanol and plated onto 1× Murashige and Skoog (MS) basal salt medium supplemented with 1× MS vitamins, 1% (wt/vol) sucrose, 0.05% MES (wt/vol), and 0.24% (wt/vol) Phytagel. After stratification in the dark at 4 °C for 5 days, plates were transferred to an incubator (Conviron) set to long days (LD) (16L:8D at 22 °C:20 °C), with light supplied at 100 μmol·m-2·s-l by cool-white fluorescent bulbs; for geldanamycin experiments, plates were placed under long-pass yellow filters that blocks 454 nm light (67). Seedlings were scored for mutant phenotypes at day 10. Geldanamycin was purchased from Sigma Aldrich (G3381), diluted in DMSO, and used for M_1_ and M_2_ experiments. As this reagent was no longer available from Sigma at reasonable cost, another lot of geldanamycin was purchased from LC Laboratories (G-4500), diluted in DMSO, and used in the M_3_ heritability experiments. This reagent showed stronger phenotypic effects than Sigma geldanamycin and hence was used at a lower dose (0.5 μM instead of 0.6 μM)

### EMS Mutagenesis

For each genotype and treatment, 50 mg of seeds were weighed out (~2500 seeds). Seeds were sterilized by a ten-minute wash in 70% ethanol (v/v) with 0.1% Triton-X (v/v), followed by a five-minute wash with 95% ethanol (v/v) and one wash with sterile water. Seeds were then transferred to a tube containing 12 ml of 0.0005% Tween (v/v). EMS (ethyl methanesulfonate, Sigma-Aldrich, M0880) was added to a final concentration of either 0.2% (v/v) or 0.4% (v/v) in each tube. Control samples received only Tween treatment without EMS. Tubes were rotated at room temperature for 15 hours. Seeds were washed eight times for ten minutes in sterile water and then plated on indicated plant growth medium or sown on soil.

### Determining viable seeds and embryonic lethality in M_1_ progeny

After mutagenesis, M_1_ seeds were sown on soil and grown as indicated. For each genotype and treatment, one silique was collected from ~ 50 adult plants. For each silique, the number of viable and dead seeds was counted. Dead seeds are representative of individuals with lethal mutations and were used to estimate mutation burden after mutagenesis (49). We calculated the average number of embryonic lethal mutations per diploid genome (μ) as follows: if X equals the number of siliques without dead seeds, and Y equal the total number siliques observed, μ = −ln(X/Y). If μ is greater than one (64% or more of observed siliques have dead seeds; 1-X/Y = 0.64), mutagenesis is successfully saturated (49).

### Assessing the frequency of aberrant morphological phenotypes in the M_1_ and M_2_ generation

We devised a mutant scoring index to estimate frequency of mutant-like phenotypes quickly and objectively after mutagenesis. We based this index on known HSP90-responsive phenotypes, such as size-related traits or lesions on aerial tissue (29, 42, 56). In this index, 16 complex early seedling traits are categorized by severity (**Figure 2A**). Traits such severe developmental delay and lethality were weighted as more deleterious. For example, a seedling with the phenotype of severe developmental delay received a score of 3, while a seedling with epinastic cotyledons received a score of 1 (See Figure 2B **for examples**). The mutant scoring index also accounted for seedlings with multiple mutant phenotypes; these received higher index scores (See Figure 2B **for examples**).

Seedlings were grown as previously described (29, 42). Seedlings of indicated genotypes and treatments were scored blindly for the described morphological phenotypes (**Figure 2**). Seedlings were photographed with a Canon Power Shot S5 IS camera, pictures were edited using Adobe Photoshop CC 2017. Edits addressed image size, brightness, and contrast and did not alter depiction of seedling phenotypes.

### M_3_ heritability experiments

65 M_2_ individuals with aberrant, deformed aerial organ phenotypes, grown on plates containing either 0.5 μM GdA or DMSO, were selected, transferred to soil, and propagated to generate M_3_ seeds (see **S3 Table for parental phenotypes**). For the selected seedlings, ~50% survived the transfer and produced progeny. The resulting M_3_ progeny, designated either DM (progeny of M_2_ plant with selected phenotype grown on DMSO medium) or GM (progeny of M_2_ plant with selected phenotype grown on GdA medium), was assayed for 16 early-seedling phenotypes in the presence of either 0.5 μM GdA or DMSO (GdA solvent, mock treatment). For phenotyping, M_3_ lines were grown as indicated and scored blindly for the 16 morphological phenotypes (**Figure 2**); blind scoring relied on scrambled line identifiers. Seedlings were photographed with a Canon Power Shot S5 IS camera and were edited using Adobe Photoshop CC 2017.

A given trait in a given M_3_ line was considered responsive to HSP90 perturbation if *X*_GdA, M3_ − *X*_DMSO, M3_ ≥ 10% (positive) or if *X*_DMSO, M3_ − *X*_GdA, M3_ ≤ 10% (negative) with *X*= (%) phenotype frequency of affected seedlings. A trait was considered unresponsive to HSP90 perturbation if *X*_GdA, M3_ − *X*_DMSO, M3_ ≈ 0. Further, to account for effects of HSP90 perturbation on traits commonly observed in the wild-type Col-0 background, we considered traits with *X*_GdA, M3_ − *X*_DMSO, M3_ ≈ *X*_GdA, wild-type Col-0_ − *X*_Dmso, wild-type Col-0_ as showing no effect of newly introduced EMS mutations. Comparisons of seedling phenotype frequencies were performed with Fisher’s exact test to determine statistical significance.

### SNV calling on two unmutagenized Col-0, two unmutagenized HSP90 RNAi-Cl, six mutagenized Col-0 and six mutagenized HSP90 RNAi-Cl individual plants

Genomic DNA was extracted from one adult leaf per individual using the CTAB method. Total DNA was quantified with Qubit HS dsDNA assays according to the manufacturer’s instructions. Sequencing libraries were generated using 50 ng of input DNA using the Nextera (Illumina) sample kit according to the manufacturer’s instructions. Libraries were quality checked on the Agilent 2100 bioanalyzer using a DNA high-sensitivity chip (Agilent). The samples were sequenced by Genewiz on an Illumina HiSeq (100bp paired-end reads).

WGS reads were aligned to the TAIR10 release genome (http://www.arabidopsis.org/) using the default parameters of BWA (68). Aligned reads with mapping quality less than 35 were discarded. A VCF file was generated using SAMtools (http://samtools.sourceforge.net). Variants were further filtered for minimum coverage = 5 and minimum derived allele coverage = 2. We filtered each VCF file for unique mutations (*i.e*. unshared with any other sample within either Col-0 or HSP90 RNAi-C1 genotypes) as these were likely due to the EMS treatment. After identifying unique and private SNVs, we determined the number of total non-reference transitions and transversions for each individual. For analysis in mutagenized backgrounds, a similar filtering process was performed, where again we determined unique, unshared non-reference variants in each sequenced individual. See **S1 Table** for more information about the called variants.

### Bulk segregant sequencing analysis

We backcrossed M_3_ individuals with deformed aerial organs from the GM6 and GM9 lines to Col-0 wild-type plants used for mutagenesis. The resulting F_1_ plants were propagated to the F_2_ generation. For each backcross, F_2_ seedlings were grown on HSP90-reduced growth medium (0.5 μM GdA), seedlings with deformed aerial organs were selected and pooled (n>90). Total DNA was extracted from these pools using the CTAB method; DNA was quantified using the Qubit HS dsDNA assayed. Sequencing libraries were generated using 10 ng input DNA with the Nextera (Illumina) sample kit. Libraries were quality checked on the Agilent 2100 bioanalyzer using a DNA high-sensitivity chip (Agilent). The samples were sequenced using an Illumina NextSeq in 150bp paired-end run.

Because our Col-0 line has been bulk-propagated in our lab since 1994, wild-type Col-0 individuals contain derived SNVs with respect to the published TAIR10 reference sequence at varying frequencies. We identified these background SNVs by sequencing two pooled Col-0 individuals at ~8 x-coverage, allowing us to remove such derived SNVs from our list of EMS-generated mutations. Additional background Col-0 SNVs were identified by comparing SNVs in the deeply sequenced (>50 x-coverage each) F_2_ populations derived from GM6 and GM9. Derived SNVs found in both populations are unlikely to be due to EMS treatment.

With the pooled F_2_ plants, we are measuring derived allele frequency at likely EMS-generated SNVs, a task that requires much greater read depth. Furthermore, SNVs present in the unmutagenized parents (M_0_) relative to TAIR10 reference genome must be carefully filtered out. As we are attempting to identify regions rather than single SNVs with elevated derived allele frequency, erroneously discarding or including certain SNVs should not prevent the identification of regions containing causative SNVs.

Our filtering pipeline proceeded as follows:

First, reads from the F_2_ pools were mapped to the TAIR10 reference genome using BWA (default parameters). Mapped reads outside of the centromeres (chr1:13,698,788-15,897,560; chr2:2,450,003-5,500,000; chr3:11,298,763-14,289,014; chr4:1,800,002-5,150,000; chr5:10,999,996-13,332,770) with map quality > 20 were maintained (69).

Next, SNVs were identified in a pool of M_0_ Col-0 seedlings to identify background SNVs as well as in the two pools of F_2_ seedlings (aberrant seedlings grown on GDA-reducing media, for both the GM6 and GM9 strains). All bases with any evidence of an alternate allele were identified (derived allele frequency > 0). We excluded background SNVs (DAF>0) identified (i) in the Col-0 pool (n=1,022,369), and (2) in both the two deeply sequenced GM6 and GM9 GDA-treated F_2_ pools (n=1,093,093). There was very little overlap between these sets of background SNVs (n=78,805), indicating that the vast majority of the former set were either spuriously identified as polymorphic -- which is certainly possible given the low sequencing coverage -- or neither of the individual seeds that were mutated to give rise to the GM6 and GM9 lines contained the derived allele -- also possible, given that many of these flagged bases had relatively low derived allele frequency.

Among this filtered set of EMS-derived SNVs identified in the two F_2_ pools, we then attempted to remove SNVs erroneously arising due to sequencing error by removing SNVs with less than two reads indicating the derived allele. Reads collected typically had per-bp Phred quality score of 20 or more (error rate < 1/1,000), therefore the probability of two reads presenting an identical error is low.

We also attempted to remove SNVs erroneously arising due to mapping error by removing SNVs with unusually low x-coverage. The mean x-coverage in the GDA-treated F2 pools was high (GM6 F_2_ x-coverage=77; GM9 F_2_ x-coverage=54) and followed an approximately lognormal distribution, per base. Specifically, we required each such bp to have xcov greater than mean(log10(x-coverage)) - 2*sd(log10(x-coverage)).

For our final analysis, we examined only likely EMS mutations with derived allele frequency > 0.2.

### Statistical analysis

All statistical analyses were performed in R Version 3.2.5. Comparisons of seedling aberrant phenotype frequencies were performed with Fisher’s exact test (**S2 Table**). Comparisons of embryonic lethality were modeled with a Poisson regression using a general linear model. Student’s t-tests performed for seed viability. When appropriate, p-values were adjusted using the R function “p.adjust,” with method= “fdr.” The results of all statistical tests performed are in **S2 Table**.

## Acknowledgments

We thank Josh Cuperus, Cristina Alexandre, and Ken Jean-Baptiste and other members of the Queitsch lab for technical assistance and important conversations. We thank Stan Fields for edits and comments on the manuscript. We thank the Fields lab for access to and help with sequencing instruments. We thank UW Genome Sciences Information Technology for high performance computing resources. Finally, we thank Sandra Malone for moral support.

## Supporting Information

**S1 Fig. Analysis pipeline for SNV calling in M2 plants.** See Materials and Methods for details and S1 Table for results.

**S2 Fig. HSP90 perturbation increases penetrance of new mutations in the M_1_ generation. (A)** Schematic of experimental work flow. **(B)** Genetic perturbation of *HSP90* yields significantly increased frequency of aberrant phenotypes in M_1_ HSP90 RNAi-C1-seedlings. Seeds of the indicated genetic backgrounds were treated with 0.2% EMS, plated on growth medium, seedlings were grown for ten days under standard growth conditions and assessed for 16 early-seedling traits (see **Figure 2**). The following number of seedlings was examined: wild-type Col-0 control n=213; mutagenized Col-0 M_1_ n=948; un-mutagenized HSP90 RNAi-A1 control n=206; mutagenized HSP90 RNAi-A1 M_1_ n=1009; un-mutagenized HSP90 RNAi-C1 control n=206; mutagenized HSP90 RNAi-C1 M_1_ n=961. *p <0.05, Fisher’s exact test. Error bars represent standard error as calculated by: SE = sqrt(*p*(100-*p*)/*n*), where *p* equals the percentage of affected seedlings and *n* equals the number of observations (see **S2 Table** for all odds ratios and p-values). **(C)** Pharmacological inhibition of HSP90 reproduces the result observed in (**B**) for HSP90 RNAi-C1. Seeds, treated with either 0.2% or 0.4% EMS, were plated on media containing either DMSO or 1.0 μM GdA, grown for ten days under standard growth conditions and assessed for 16 early-seedling traits (see **Figure 2**). The following number of seedlings was examined: wild-type Col-0 control on DMSO n=144; 0.2% EMS-treated Col-0 on DMSO n=530; 0.4% EMS-treated Col-0 n=355 on DMSO; wild-type Col-0 control on 1.0 uM GdA n=36; 0.2% EMS-treated Col-0 on 1.0 uM GdA n=540; 0.4% EMS-treated Col-0 n=329. ***p-adj<1.0E-08, Fisher’s exact test.

**S3 Fig. Aberrant phenotypes are heritable and HSP90-responsive across generations in several mutagenized lines.** Frequencies of 15 aberrant phenotypes in wild-type Col-0 and mutagenized M_3_ lines in response to HSP90 perturbation (0.5 uM GdA) or mock conditions (DMSO). Black lines represent the response of wild-type Col-0 to HSP90 perturbation in three biological replicates. Shades of red denote lines with 10% (5% for frequency of aborted seedlings) increase in phenotype frequency under HSP90-reduced conditions and shades of blue denote lines with 10% decrease in phenotype frequency under HSP90-reduced conditions (5% for frequency of aborted seedlings). Gray line color denotes non-responsive lines. The following number of seedlings was examined per treatment: wild-type Col-0 n~144 in each of three replicate experiments; mutagenized M_3_ lines n~72.

**S4 Fig. Several M_3_ lines propagated from M_2_ parents with deformed aerial organs show higher frequency of this trait in response to HSP90 perturbation than Col-0 wild-type.** Heatmap of clustered odds ratios for 0.5 μM GdA-treated wild-type Col-0 seedlings vs. 0.5 μM GdA-treated seedlings of indicated M_3_ lines as calculated for each of the 16 early-seedling phenotypes. Black asterisk (*), p-adj<0.05, Fisher’s exact test, significantly greater trait frequency in indicated M_3_ lines compared to wild-type Col-0 seedlings in response to HSP90 perturbation.

**S5 Fig. Bulk segregant analysis identifies candidate locus associated with HSP90-responsive trait. (A)** Schematic of experimental work flow indicating provenance of examined seedling populations. **(B)** Image of the selected M_2_ parent GM9, showing the deformed aerial organ phenotype that was successfully propagated to the M_3_ generation. Scale bar equals 5mm. **(C)** The GM9 M_3_ line showed increased frequency of seedlings with deformed aerial organs compared to wild-type Col-0. A GM9 M_3_ individual with deformed aerial organs was then backcrossed to Col-0. The resulting F_2_ population showed significantly increased frequency of the deformed organs trait when HSP90 was inhibited with GdA. Errors bars represent standard error of the mean across two biological replicates. *p-adj<0.05, **p-adj<1.0E-04, ***p-adj<1.0E-14, Fisher’s exact test. **(D)** Bulk segregant analysis of GM9 F_2_ populations identifies a candidate locus associated with the HSP90-responsive trait deformed aerial organs. F_2_ individuals displaying on HSP90-reduced conditions (n=117) were sequenced. Derived allele frequencies (y-axis) were estimated as the fraction of reads supporting the derived allele divided by the number of reads mapping to a given locus (see **Materials and Methods**). Dark gray box indicates the centromere and light gray box represents pericentromeric regions (chr1:12,000,000-18,000,000; chr2:1,000,000-7,500,000; chr3:11,000,00-17,000,000; chr4:1,800,000-7,500,000; chr5:10,000,000-17,000,000). Dashed red line represents derived allele frequency for the lowest observed derived allele frequency for the six candidate mutations (**S4 Table**). Dashed black line represents expected derived allele frequency of 0.5. The candidate locus contains six transition SNVs (**S4 Table**).

**S6 Fig. Bulk segregant analysis data of backcrossed GM6 and GM9 F_2_ populations.** Allele frequency estimations at likely EMS-derived SNVs in GM6 **(A)** and GM9 **(B)** backcrossed F_2_ populations. F_2_ individuals displaying this trait on HSP90-reduced conditions (GDA) were sequenced. Derived allele frequency estimations across all five chromosomes are shown (bp; x-axis). Dark gray boxes indicate centromeres(1) and light gray boxes represent pericentromeric regions (chr1:12,000,000-18,000,000; chr2:1,000,000-7,500,000; chr3:11,000,00-17,000,000; chr4:1,800,000-7,500,000; chr5:10,000,000-17,000,000). Derived allele frequencies (y-axis) were estimated as described in the **Materials and Methods**. Dashed red line represents derived allele frequency for the lowest observed derived allele frequency for the six candidate mutations. Dashed black line represents expected derived allele frequency of 0.5. The candidate locus contains six transition SNVs (**S4 Table**).

**S7 Fig. Derived allele frequencies for 100 simulations.** Simulation, described fully in the Materials and Methods section, assumes a given number of SNVs (*n*), recombining length of chromosome (*L*), and x-coverage (*x*), each based on actual data. The final simulation is shown in bold to illustrate typical levels of fluctuation in derived allele frequency for one simulation. Note that while derived allele frequency at simulated SNVs at times exceeds 0.65 (red dashed line), none of the 100 simulations contained five or more consecutive SNVs with this frequency.

S1 Table. Number of SNVs in mutated and unmutated plants.

S2 Table. All p-values.

S3 Table. Parental M2 phenotypes.

S4 Table. Candidate SNVs with elevated DAF.

